# Damage Repair Versus Aging in Biofilms

**DOI:** 10.1101/2020.01.08.899740

**Authors:** Robyn J. Wright, Robert J. Clegg, Timothy L. R. Coker, Jan-Ulrich Kreft

**Author notes:** Address correspondence to Robyn J. Wright,;. Author contributions: JUK and RJC initially designed the study with later input from RJW and TLRC. All simulations and analyses shown were carried out by RJW with guidance from RJC and JUK. TLRC performed preliminary simulation experiments to choose many of the simulation parameters. RJW and JUK wrote the first draft of the manuscript and all authors contributed to revisions. The authors declare no conflicts of interest.

## Abstract

The extent of senescence due to damage accumulation (or aging) is evidently evolvable as it varies hugely between species and is not universal, suggesting that its fitness advantages depend on life history and environment. In contrast, repair of damage is present in all organisms studied. Repair and segregation of damage have not always been considered as alternatives, despite the fundamental trade-off between investing resources into repair or growth. For unicellular organisms, unrepaired damage could be divided asymmetrically between daughter cells, leading to aging of one and rejuvenation of the other. Repair of unicells has been shown to be advantageous in well-mixed environments such as chemostats. However, most microorganisms live in spatially structured systems such as biofilms with gradients of environmental conditions and cellular physiology as well as clonal population structure. We asked whether this clonal structure might favor aging by damage segregation as this can be seen as a division of labor strategy, akin to the germline soma division in multicellular organisms. We used an individual-based model with a newly developed adaptive repair strategy where cells respond to their current intracellular damage levels by investing into repair machinery accordingly. We found that the new adaptive repair strategy was advantageous whenever efficient and optimal, both in biofilms and chemostats. Thus, biofilms do not favor a germline soma-like division of labor between daughter cells in terms of damage segregation. We suggest that damage segregation is only beneficial when active and effective, extrinsic mortality is high and a degree of multicellularity is present.

**IMPORTANCE:** Damage is an inevitable consequence of life, leading to a trade-off between allocating resources into damage repair or into growth whilst allowing aging, *i.e.*, segregation of damage upon cell division. Few studies considered repair as an alternative to aging. Moreover, all previous studies merely considered well-mixed environments, although the vast majority of unicellular organisms live in spatially structured environments, exemplified by biofilms, and fitness advantages in well-mixed systems often turn into disadvantages in spatially structured systems. We compared the fitness consequences of aging versus damage repair in biofilms with an individual-based model implementing an adaptive repair mechanism based on sensing damage. We found that aging is not beneficial. Instead, it is useful as a stress response to deal with damage that failed to be repaired when (i) clearly asymmetric cell division is feasible; (ii) extrinsic mortality is high; and (iii) a degree of multicellularity is present.

## INTRODUCTION

Senescence is all around us, yet it is not obvious why it has evolved in many taxa as it would appear to be detrimental to the fitness of individuals. Importantly, the extent of senescence, manifesting in decreasing fecundity and/or increasing mortality with age, is clearly evolvable as it varies hugely between species and is *not* universal. For example, several taxa of simple multicellular organisms can fully regenerate and for several taxa of complex multicellular organisms, fecundity does not simply decrease with age and/or mortality does not simply increase with age (1–3). An evolutionary explanation for the various extents of senescence present in different organisms is challenging, particularly for unicellular organisms that divide apparently symmetrically, such as most prokaryotes and some eukaryotic unicells, in contrast to multicellular animals with a clear division of labor between germline and soma (4, 5). However, the first single-cell study of division asymmetry in *Escherichia coli* highlighted that morphological symmetry does not exclude functional asymmetry as daughter cells inheriting the old cell pole were shown to grow a little slower than the mother cell while the new cell pole daughters grew a little faster (6). Ironically, *Caulobacter crescentus*, the bacterium first studied in terms of aging (7), as it has substantial morphological and functional asymmetry in cell division, has been shown in more recent high-throughput microfluidic studies to maintain a constant growth rate over cell divisions under benign conditions (8) and to divide protein aggregates symmetrically between mother and daughter cells (9).

Following the first single-cell studies that suggested the existence of aging in unicellular prokaryotes (7, 10, 11) and unicellular eukaryotes (12), there has been a gold rush of studies eager to demonstrate aging in further unicells, such as bacteria (13–15) and eukaryotic algae (16–19). However, the loss of fecundity (10) or increase of mortality (20) with age, demonstrated in some of these unicells, are rather small effects compared with the resource limitations of growth and high external mortality in most environments. The effects are also much smaller than in the budding yeast, which has long been known to have a limited replicative lifespan (21), supported by several recent high-throughput single-cell studies (22–28). However, it may be misleading to regard the budding yeast as a unicellular organism as wild relatives are capable of dimorphic growth (29) and domesticated strains rapidly evolve multicellularity (30, 31). Crucially, a number of recent experimental results have led to a reinterpretation of aging, primarily in the sense of segregating protein aggregates, as a stress response rather than an evolved characteristic of growth under benign conditions (9, 20, 32–38).

In concert with the gold rush for experimental evidence for aging, there has also been one of mathematical modelling studies eager to find evolutionary advantages of aging. Some of these models did not consider extrinsic mortality (39–41), although it favors rapid and early reproduction and thus tilts the evolutionary trade-off towards investment of resources into growth and reproduction, rather than maintenance and repair (1, 42). Some also did not consider repair as an alternative (40, 41, 43, 44) or did not consider the cost of repair (39).

Repair is present in all organisms studied and evidence for the evolution of mechanisms that repair damage, such as misfolded and aggregated proteins (9), is beyond doubt. Of the few models that consider both extrinsic mortality and costly repair, the model of Ackermann *et al.* (2007) (45) was the first. It found that asymmetric damage segregation outperformed repair. In contrast, our previous study, Clegg *et al.* (2014) (46), found that repair was always beneficial, while damage segregation was beneficial only in addition to repair and if three conditions were fulfilled simultaneously: (i) damage is toxic, (ii) damage accumulates at a high rate and (iii) repair is inefficient. The reason for this discrepancy could be pinned down. In Ackermann *et al.* (2007) (45), rather than growing, cells divide at fixed time intervals, while in Clegg *et al.* (2014) (46), cells grow by consuming substrate and divide when they have reached a threshold size. This enabled both an immediate benefit of repair (increased growth rate of a less damaged cell leading to earlier division) and an immediate cost of repair (diverting resources away from growth to repair machinery). Overall, this made repair advantageous.

More recent models that also consider repair have come to similar conclusions (36, 47, 48). Vedel *et al.* (2016) (36), in particular, has advanced our understanding with an experimentally validated model that considers the fitness of the whole lineage. This helped to identify a positive feedback loop, shifting growing lineages towards less damaged cells and explaining how higher stress levels lead to higher damage accumulation which in turn leads to higher damage segregation.

None of these studies considered the fitness effects of aging and repair in spatially structured environments such as biofilms, although biofilms are prevalent in nature and important for ecosystem function. For humans, they have many advantages in biotechnology but also cause big problems in industry and health. Biofilms are heterogeneous in both time and space (49) and cells that are growing within them are therefore exposed to varying, and often limiting, nutrient regimes (50). This leads to gradients in growth rate and the presence of an active layer in biofilms, where active growth only occurs close to the boundary of the biofilm, due to slow nutrient diffusion (51). Growth within biofilms has also been shown to confer tolerance to damage-inducing agents, such as antibiotics (49, 52–54) and UV radiation (55, 56), and it is therefore likely that these gradients of growth rate and stress could make the evolutionary benefits of aging and repair different from spatially uniform environments, such as chemostats. Moreover, biofilms have a clonal population structure unless the cells remain motile (57–59). This can have strong effects on the evolution of division of labor (60). Damage segregation can be seen as a division of labor akin to the germline soma differentiation in multicellular animals (61). Thus, we hypothesized that biofilms might favor damage segregation over repair.

To test this hypothesis, we developed a new individual-based model with adaptive repair, whereby cells were able to sense and respond to their current intracellular damage levels. This enables an appropriate response to gradients of stress and damage in biofilms. We found that adaptive repair rather than damage segregation or a fixed rate of repair was the optimal strategy for unicells growing in a biofilm, but only when the rate of damage accumulation was proportional to the cells’ specific growth rate.

## RESULTS

### Characteristics of adaptive repair

We developed a new repair strategy where allocation of newly synthesized protein into repair machinery, denoted by *β̂*, rather than growth machinery, depends on the current level of damage in the cell and compared this with previous strategies (Fig. 1). The idea was that the cell can sense and appropriately respond to its damage level. This adaptive repair is more appropriate than a fixed repair strategy when rates of growth and damage accumulation vary in time or space, *e.g.* in fluctuating or spatially structured environments, such as biofilms. Adaptive repair is therefore meant to replace the previous strategy of a fixed allocation into repair machinery at an optimal level, *β*, which is appropriate for constant or chemostat environments, where growth and damage accumulation rates are in a steady state, like in our previous study, Clegg *et al.* (2014) (46). There, we estimated the optimal, fixed investment into repair (here simply referred to as fixed repair) by examining the mean specific growth rates of cells with different investments into repair and found that for a damage accumulation rate of 0.1 h^-1^, an investment into repair of *β* = 0.07 was optimal for both asymmetric and symmetric strategies (for the case where damage is toxic, that we focus on here). The purpose of this section is to examine the consequences of the new adaptive repair strategy and to test whether it is equivalent to the previous optimal repair strategy in steady state environments.

**FIG 1.**
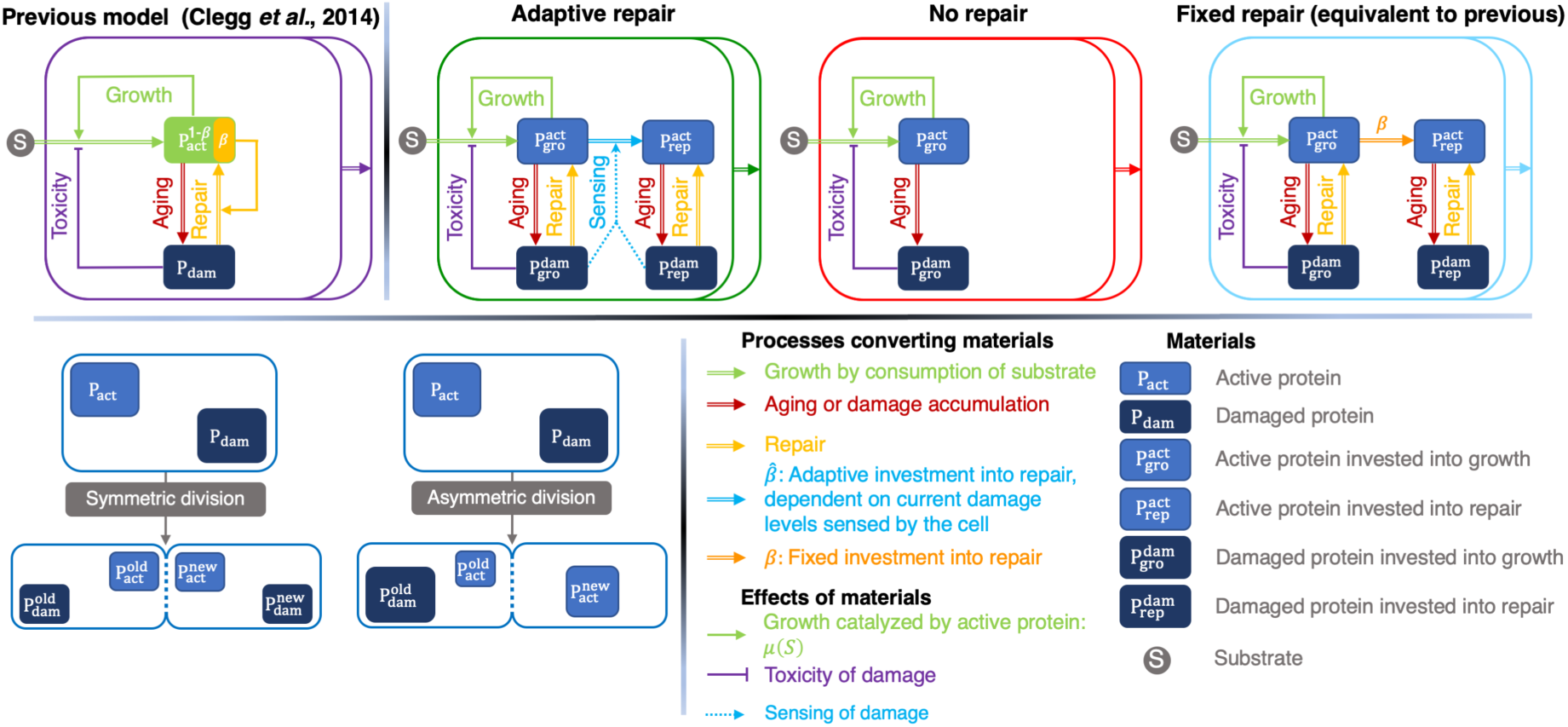
General schematic of division and repair strategies used in this study, giving rise to 6 combinations: (i) symmetric division with no repair (NR) (ii) symmetric division with fixed repair (FR); (iii) symmetric division with adaptive repair (AR); (iv) asymmetric division (damage segregation) without repair (DS); (v) asymmetric division with fixed repair (DSFR); and (vi) asymmetric division with adaptive repair (DSAR).

### Consequences of adaptive repair on the fraction of repair protein in single cells

The adaptive repair strategy leads to an allocation into repair that responds to current levels of damage and therefore lags behind the ideal level of repair machinery, unless damage levels reach a steady state due to symmetric division (Fig. S1). Since asymmetric division causes sudden changes in damage levels, the current investment into repair tracks the changing damage levels (Fig. 2A). Damage levels then change as a result of repair, which in turn changes allocation into repair. As a result, the level of repair machinery never reaches ideal levels in asymmetrically dividing cells, in contrast to symmetrically dividing cells (Fig. 2A and S1). To approach ideal levels of repair machinery more quickly, a high turnover of repair protein would be required. Since cellular proteins that are not involved in regulation have a long half-life of more than one generation (62), it seems more realistic to assume that repair protein does not turn over; turnover would also be costly and unnecessary. Investment into repair varies greatly over the cell cycle for cells with asymmetric segregation of damage but is always at approximately the same level immediately before division. The investment into repair immediately before division is approximately the same as the fixed optimal investment into repair.

**FIG 2.**
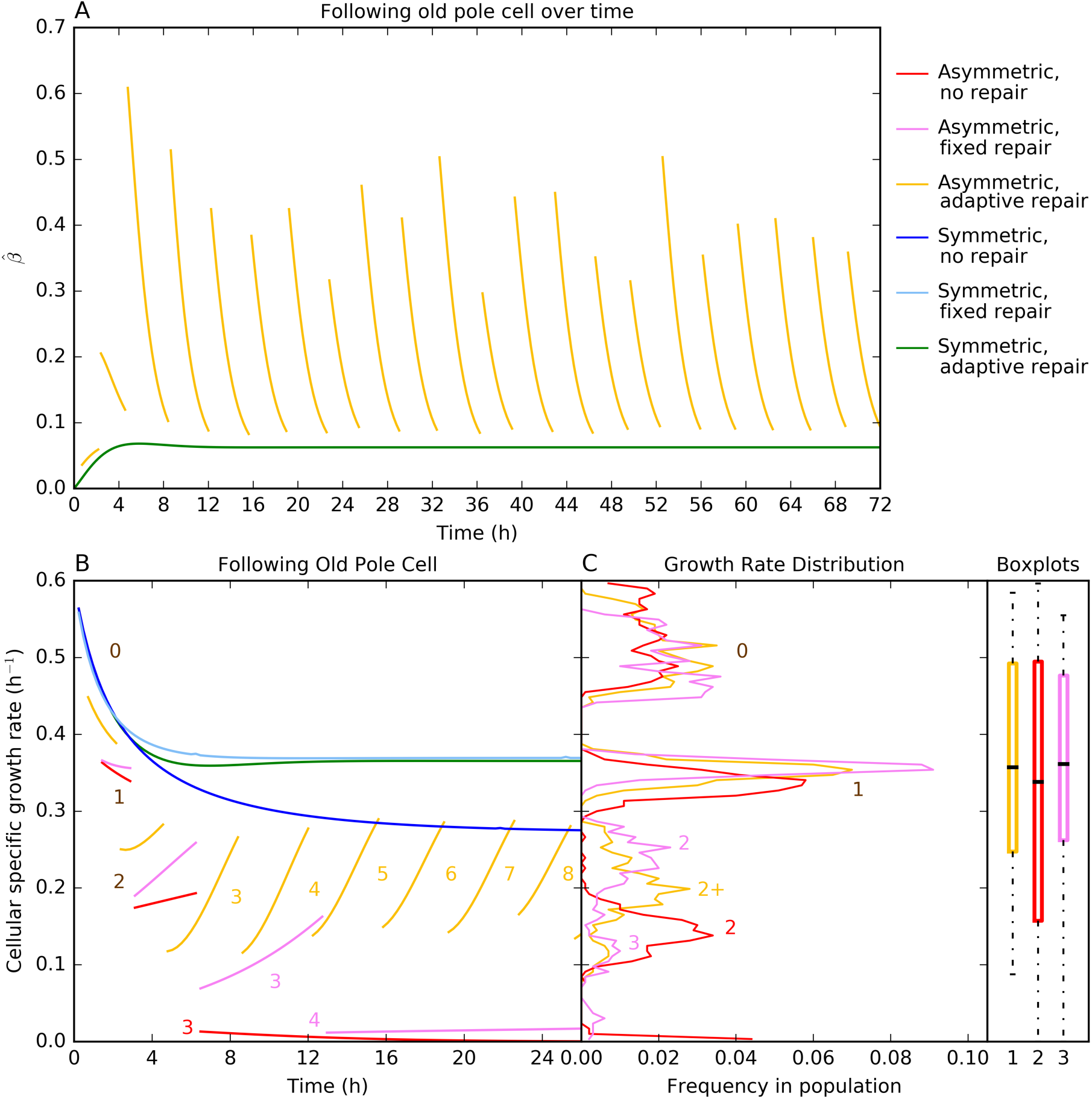
Characteristics of strategies in a constant environment (no competition). **(A)** Investment into repair (*β̂* on the y axis) for the new adaptive repair strategy following the old pole cell over many divisions. In asymmetric divisions, the old pole cell inherits all damage, leading to a jump in allocation into repair following division and then decreasing steadily until the next division. In symmetric divisions, damage, and therefore investment into repair, reaches a steady state. **(B)** Specific growth rate of a single cell over consecutive cell divisions. Numbers in the panel label generations and each generation is shown with a new line (for asymmetric strategies). The specific growth rate of an asymmetrically dividing cell with no repair (red, *β* = 0) drops quickly to zero. For a cell with fixed optimal repair (magenta, *β* = 0.07), it decreases more slowly over time but also reaches zero. For a cell with adaptive repair (yellow, *β̂* variable), it decreases only initially towards a see saw pattern, as in (A). Specific growth rates do not change at division for symmetric strategies (blue, cyan, and green) and there is no difference between daughter cells. Symmetric strategies show an initial decrease in specific growth rate before reaching a steady state, with similar values for fixed and adaptive repair and lower without repair. **(C)** Distribution of specific growth rates in populations at steady state (snapshot taken at 100 days) for asymmetrically dividing cells. Specific growth rates of cells with adaptive repair are between those with fixed repair and those without repair. The medians and inter-quartile ranges for adaptive and fixed repair are close and higher than for symmetrically dividing cells. Data are reproduced with permission from Fig. 4A,B in (46), with the new adaptive repair strategy added. Specific growth rate was 0.6 h^-1^ and aging rate was 0.22 and dependent upon specific growth rate for all strategies. Replicate simulations are similar, see Fig. S2.

### Consequences of adaptive repair on growth rates of individual cells

For asymmetric strategies, cells with adaptive repair maintained the highest specific growth rate across consecutive divisions from approximately generation three onwards (Fig. 2B). Without any repair, specific growth rate declined rapidly towards zero. With fixed repair, specific growth rates likewise declined towards zero, albeit more slowly. For symmetric strategies, specific growth rates were similar for all strategies while damage levels were low. Later, cells with repair maintained substantially higher specific growth rates than cells without repair (Fig. 2B).

### Consequences of adaptive repair on the population level

A comparison of age and total biomass for each cell in a population shows the distribution of the levels of damage and how close the cells are to division (division is triggered when cells reach total biomass threshold; Fig. S3). For asymmetric strategies, adaptive repairers did not reach the high damage levels (old ages) of other strategies. For symmetric strategies, adaptive repairers were marginally older than fixed repairers, but much younger than cells without repair. Growth rates in populations of asymmetrically dividing cells had a multimodal distribution with marked differences between generations (Fig. 2C). There were fewer cells with very high and very low specific growth rates in populations using either fixed or adaptive repair. The growth rate distribution of the new adaptive repair strategy was in between the fixed and no repair strategies. The medians for each population confirm that the population specific growth rates were highest for cells using the fixed repair strategy, though the adaptive repair strategy was only slightly lower and showed less variation between cells (Fig. 2C). Symmetrically dividing cells all had the same specific growth rate as the single cells in Fig. 2B with the fixed repairers having a slight specific growth rate advantage again. In summary, the new adaptive repair strategy led to growth rates that were very similar to fixed repair at the population level, but on the individual level, there were fewer old cells.

### Competitions of aging strategies in constant and chemostat environments

Competitions are unambiguous and unbiased ways to measure fitness holistically. In the constant environment, cells were removed at random, which modelled extrinsic mortality, and the strategy that was left at the end had won. In the chemostat environment, cells were likewise removed randomly, but they also competed for the substrate that entered the environment with a given rate. This means that lineages that produced fewer offspring per time at the current concentration of substrate were washed out. In other words, cells with the highest specific growth rate (as dependent on substrate concentration) will emerge as the winner (on average, as removal is stochastic). The damage segregation without repair strategy (DS) quickly lost against either repair strategies (FR and AR) in both environments (Fig. 3). The winner took much longer to emerge between the two repair strategies and there were large fluctuations in the biomass ratios over the course of the simulations. In the constant environment, fixed repairers (FR) had an advantage, whereas in the chemostat, the adaptive repairers (AR) won in the end. Hence, the new adaptive repair strategy was slightly fitter than the fixed repair strategy in those natural environments that are better approximated by chemostats than constant environments, such as systems that are mixed on a reasonably short time scale and that receive resource inputs and experience removal of biomass by various means. However, more environments are spatially structured and are therefore better modelled by biofilms, so we turn to these in the next section.

**FIG 3.**
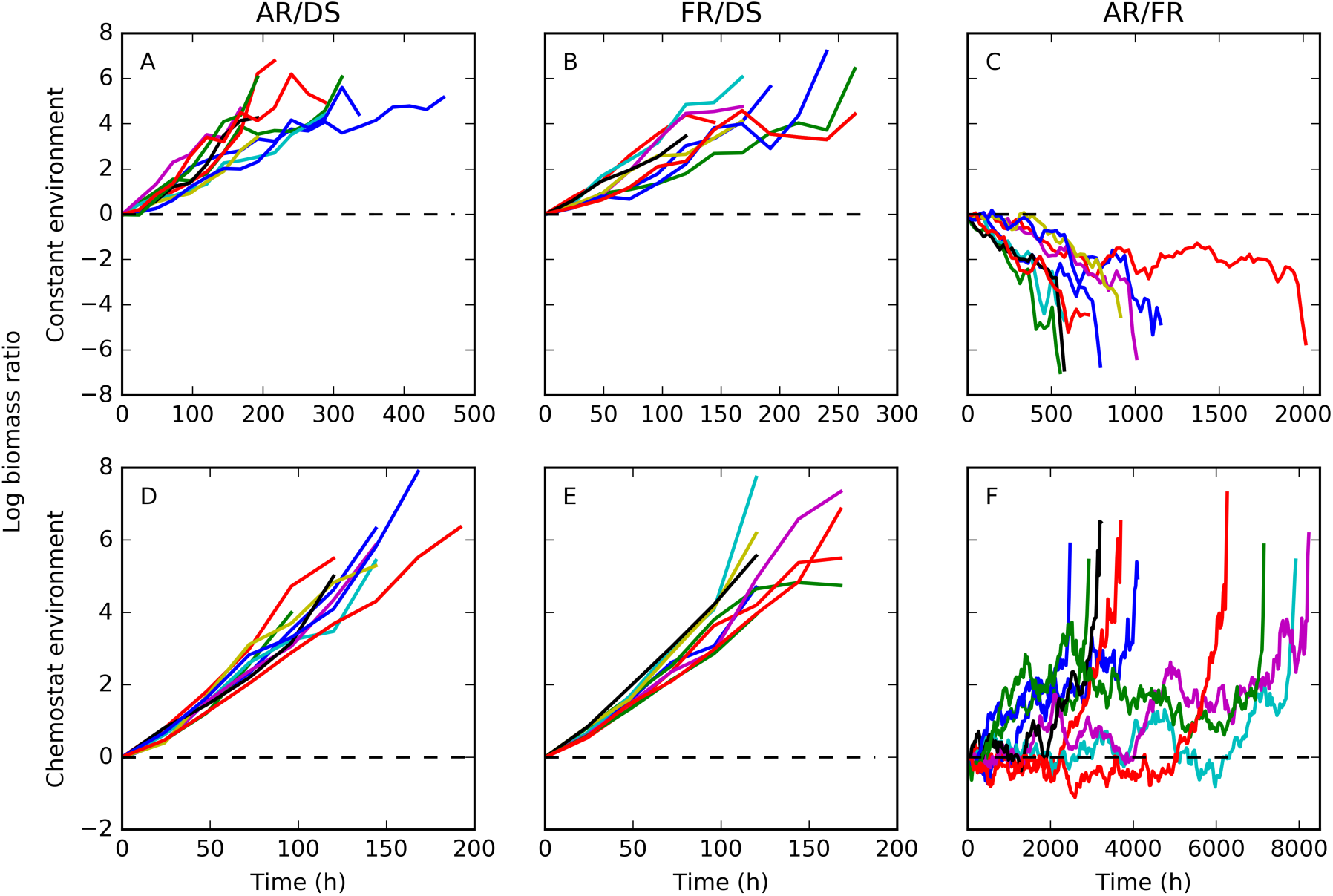
Pairwise competitions of aging strategies in constant and chemostat environments: Adaptive Repair without damage segregation (AR), Fixed optimal Repair without damage segregation (FR), Damage Segregation without repair (DS). Both repair strategies were substantially fitter than DS in either environment (A,B,D,E; *n*=10; proportion test, *p*=0.00195 for all). The two repair strategies were closer in fitness, as seen in Fig. 2B, and it took more than an order of magnitude longer before the final outcome was clear. FR was slightly fitter than AR in the constant environment (C; *n*=10; proportion test, *p*=0.00195), as expected from Fig. 2B, but not in the chemostat environment (F; *n*=10; proportion test, *p*=0.00195). Maximum specific growth rate was 0.6 h^-1^, and aging rate was 0.22 h^-1^ and dependent upon specific growth rate for all strategies.

### Generating realistic biofilm structures in the absence of damage accumulation

We first identified which parameter set would give rise to typical, rough biofilm structures with ‘finger’ formation (63–65), rather than flat biofilms, so that we could then study aging in biofilms using realistic biofilm structures (Fig. S4 and File S1). These simulations were without aging or repair. The substrate concentration in the bulk liquid was varied in order to change the dimensionless group *δ*^2^that quantifies the extent to which biofilm growth is intrinsically limited (growth-limited regime, high *δ*^2^) or limited extrinsically by diffusional mass transport into the biofilm (transport-limited regime, low *δ*^2^; see Supplementary Materials and Methods for more information on *δ*^2^). Since biofilm roughness at the end of the simulations was not significantly different between the three *δ*^2^regimes tested, we decided to continue our simulations with only one value for *δ*^2^; the intermediate value where *δ*^2^= 0.0069.

### Biofilm simulations with constant damage accumulation rate

For the biofilm simulations with constant damage accumulation rate, we compared the asymmetric damage segregation without repair (DS) and symmetric damage segregation with adaptive (AR) or fixed (FR) repair.

Repair of damaged material is assumed to require resources, *e.g.*, energy and some new material to replace the damaged parts of the old material. These resources are assumed to be supplied by endogenous metabolism of cellular material rather than the substrate, as the latter is not always available. As a result, converting damaged material into undamaged, active material comes at a loss of biomass (we assume a loss of 20% for reasons given in Clegg *et al.* (2014) (46)). This loss leads to shrinking of cells (since the density of cells is assumed to be constant), unless cells grow sufficiently fast to compensate, which is not the case in the lower layers of a biofilm. Shrinking does not affect fitness in constant and chemostat environments as only the numbers of organisms matters, but the shrinking of cells of the adaptive repair strategy had profound effects on biofilm structure and reduced the fitness of this strategy in biofilms (Fig. 4). In these competitions, the winner depended upon the initial cell density.

**FIG 4.**
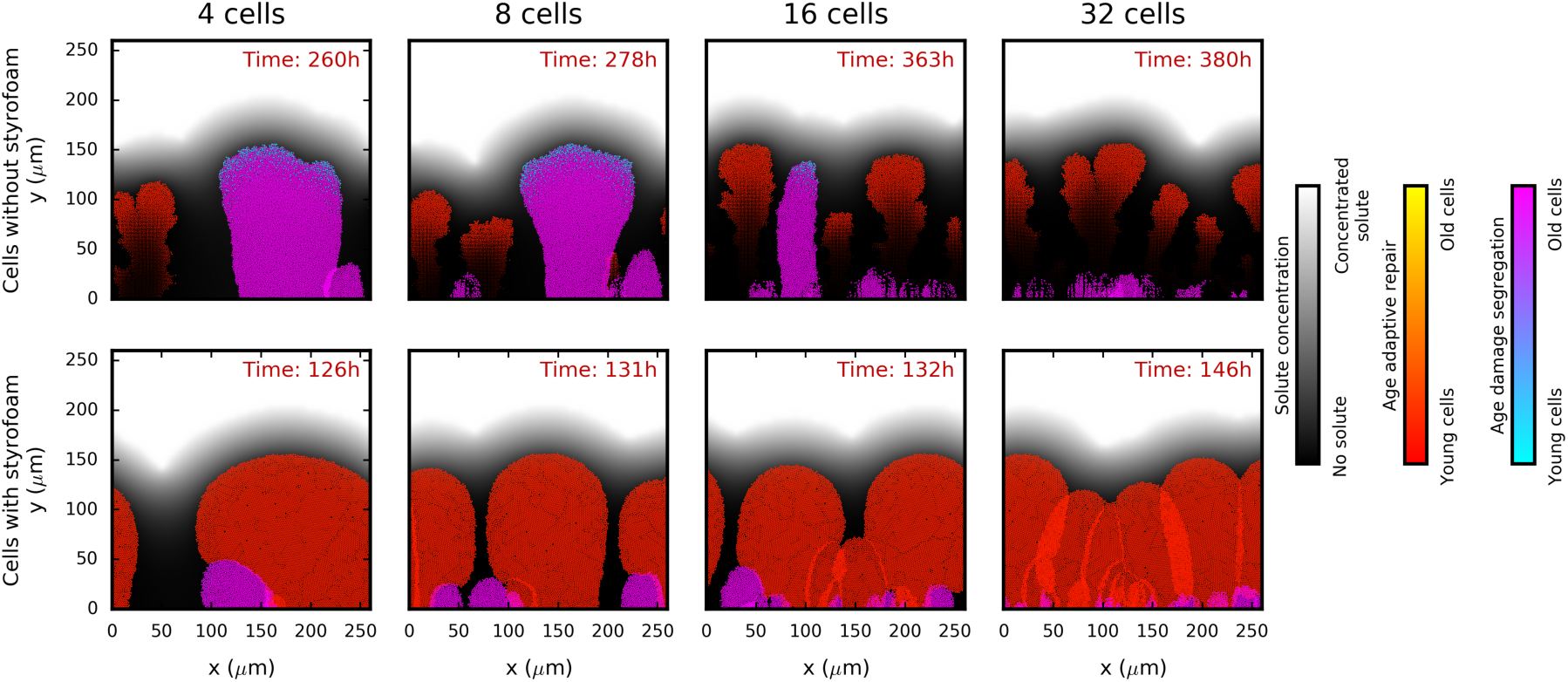
Damage segregation (DS, cold colors) *versus* adaptive repair (AR, warm colors) strategies in biofilms. Adaptive repairers are strongly affected by shrinking (top row) as they are much fitter when assumed not to shrink, *i.e.*, when the material lost due to repair is replaced with inert and massless material of the same volume (‘styrofoam’, bottom row). Initial cell density increases from left to right (4, 8, 16 or 32 cells). Biofilm structures shown have all reached a height of 154 µm. Cells are colored by age with different color gradients for each species. Maximum specific growth rate was 1.2 h^-1^ and damage accumulation rate was 0.1 h^-1^ (not proportional to specific growth rate) for all. Results of replicate simulations, simulations which competed DS and AR against the fixed optimal repair strategy (FR) and controls are shown in File S1. See Fig. S5 for time courses of log biomass ratios. For simulations without styrofoam, DS tended to be fitter than AR at initial densities of 4 and 8 cells, but AR was fitter at the higher initial densities of 16 and 32 cells (top row). For competitions between DS and FR, FR was fitter at densities of 8, 16 or 32 cells (File S1 and Fig. S5). In the simulations with ‘styrofoam’, AR and FR were always fitter than DS (bottom row; File S1 and Fig. S5). For competitions between AR and FR, FR was fitter in simulations without ‘styrofoam’, while AR was fitter in simulations with ‘styrofoam’. Control simulations that competed two cells of the same strategy always led to no clear winner.

When the initial cell density was low, it appears that the winner depends more upon the initial, random placement; cells of any strategy that happen to be placed furthest away from the other strategies tend to win. When the initial cell density is higher, the winner is more dependent upon the fitness of that strategy. Because cells of the adaptive repair strategy shrink so much more than the other strategies, this effect is much greater for them.

How much of the disadvantage of adaptive repairers was due to shrinking can be seen by comparing simulations with shrinking cells to identical simulations where, for the sake of comparison, the lost material is assumed to continue to take up volume (*i.e.*, the lost material has no mass but keeps its original volume, dubbed ‘styrofoam’). In the simulations without styrofoam, AR was only fitter than DS at initial densities of above 8 cells, FR was fitter than DS at initial densities above 4 cells and FR was fitter than AR in all simulations. In the simulations with styrofoam the results were much clearer; AR and FR are fitter than damage segregating cells and AR was fitter than FR, regardless of the cell density at the beginning of the simulation (Figs. 4 and S5).

The results caused by shrinking were thought to be unrealistic. Cells have not been observed to shrink considerably, unless they have been starved for long periods of time, and biofilm structures such as in Fig. 4 with a vanishing base due to endogenous metabolism have not, to our knowledge, been observed, and would be mechanically unstable in the presence of shear (66). Shrinking must therefore be either very limited in real cells, or the assumption that non-growing cells accumulate damage at the same rate as rapidly growing cells must be wrong (which would only cause the non-growing cells to shrink as the growing cells can make up the lost volume). We decided to avoid this unrealistic shrinking by assuming that cells that do not grow also do not accumulate damage. The damage accumulation rate was therefore assumed to be proportional to cellular specific growth rate (and was matched to the previous damage accumulation rate; Fig. S6), to allow for the very low rates of growth of cells below the active layer of the biofilm (the active layer is shown in Fig. 5). This means that shrinking is not abolished, but that slowly growing cells will accumulate damage and shrink at a lower rate as repair, leading to shrinking, is less necessary. See Materials and Methods for further explanation of this proportional damage accumulation rate. We therefore continued with a damage accumulation rate of 0.22 that is dependent upon specific growth rate and also applied this to the earlier constant and chemostat environments.

**FIG 5.**
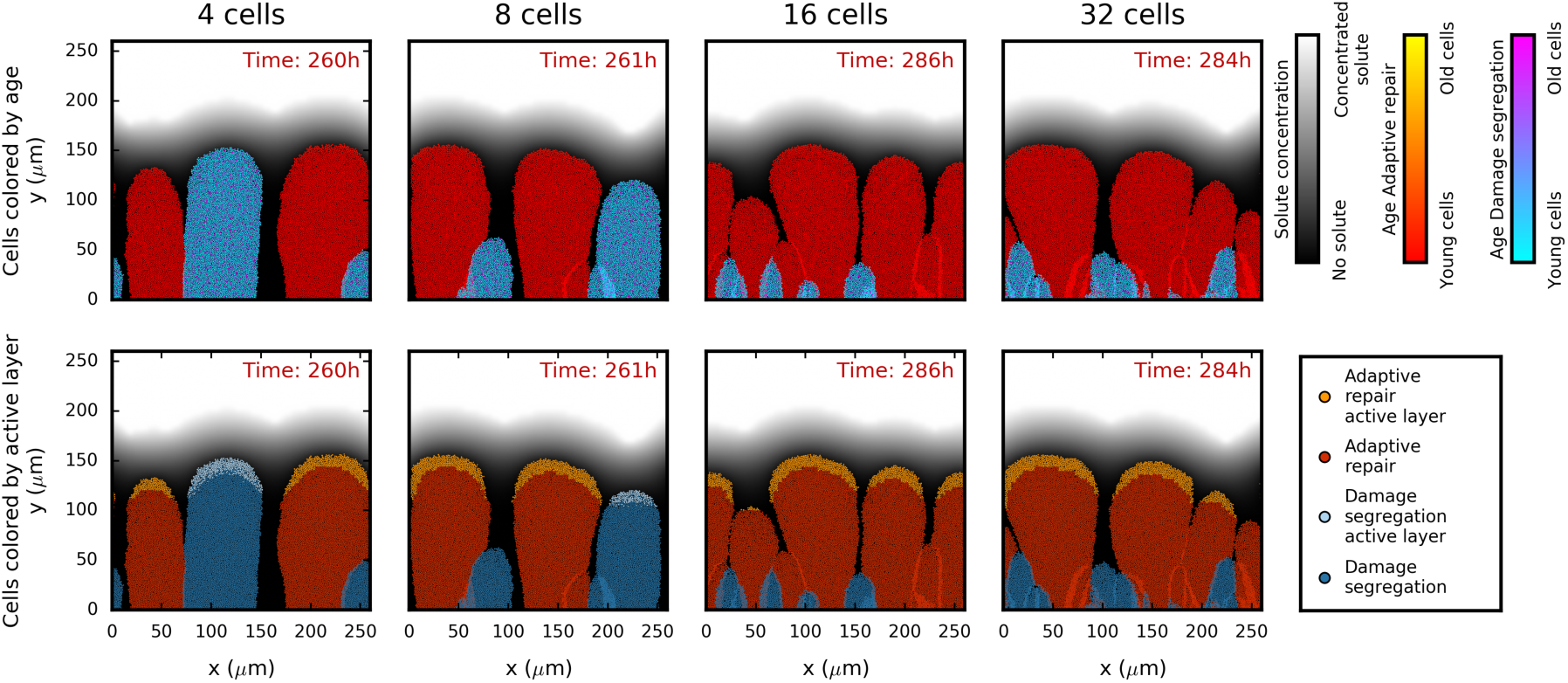
Damage segregation *versus* adaptive repair competitions in biofilms where damage accumulation rate is proportional to specific growth rate. From left to right, initial cell density increases (4, 8, 16 and 32 cells). Adaptive repair becomes more advantageous the greater the initial cell density. In the top row, cells are colored by age, while in the bottom row, cells are colored brighter if they are in the active layer of rapidly growing cells, defined as cells with a specific growth rate within 5% of the highest at this time point. Biofilm structures shown have all reached a height of 154 µm. Maximum specific growth rate was 1.2 h^-1^ and damage accumulation rate was set at 0.22 h^-1^. Repeat simulations are shown in File S1 and control competitions are shown alongside this figure in Fig. S7.

AR performed better than FR in the biofilms with styrofoam (Fig. 4), and we therefore focus on the two main alternative strategies, AR and DS, in the following section.

### Biofilm simulations where the damage accumulation rate is proportional to specific growth rate

When the damage accumulation rate was proportional to the specific growth rate, AR was more competitive than DS (Figs. 5 and 6; Fig. S7 also shows the controls). The higher the initial cell density, the stronger the advantage of AR and the earlier that they won. At the highest initial cell density (32 cells), AR won in all 50 replicate simulations (*p*=0.00). At the lowest cell density (4 cells), they won in the majority of the simulations (31 *vs* 19) but this difference was not statistically significant (*p*=0.119). The advantage of adaptive repair became statistically significant at initial cell densities of eight or higher (Table S1).

**FIG 6.**
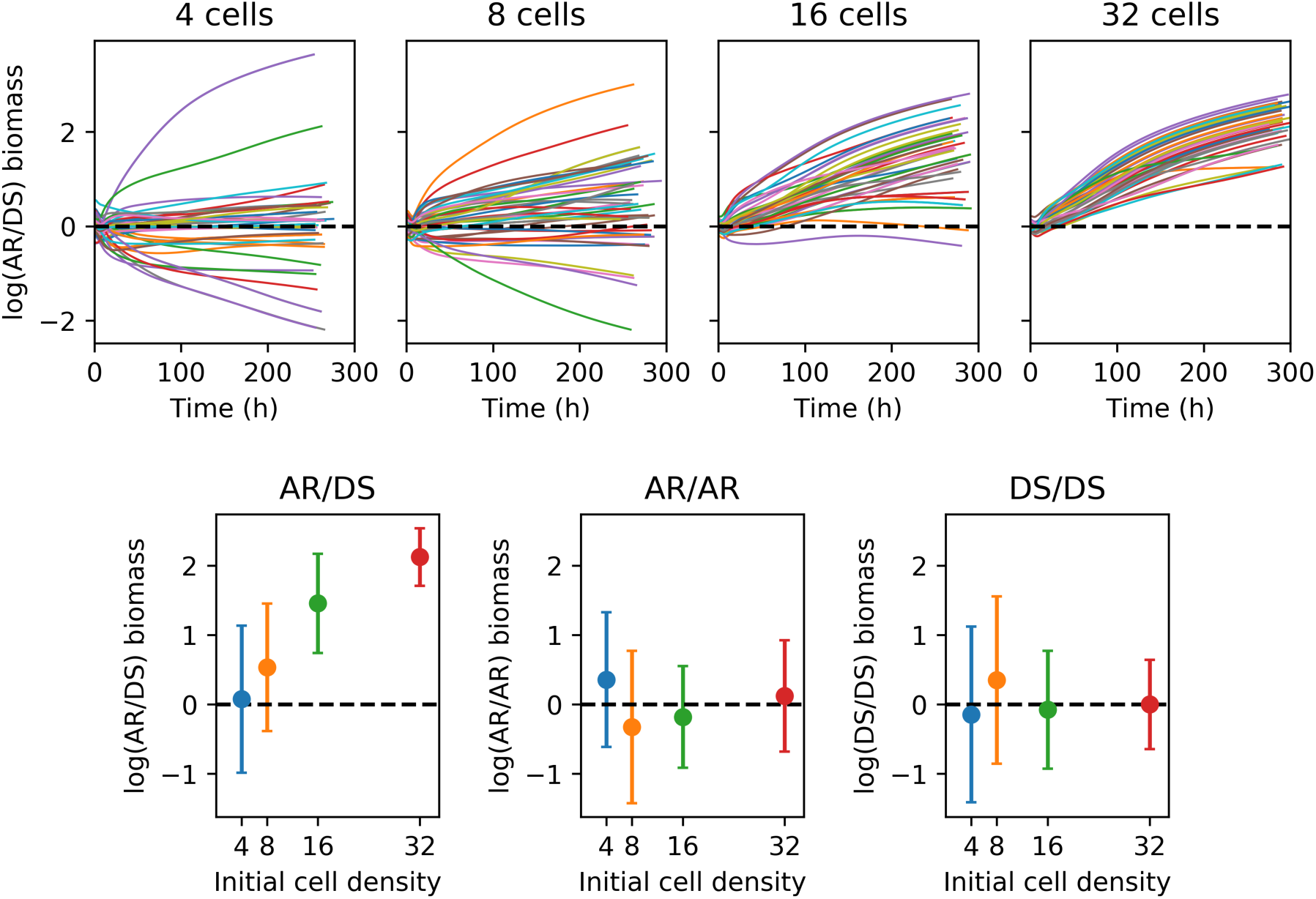
Time courses of log biomass ratios for 50 replicate biofilm competitions between adaptive repair (AR) and damage segregation (DS) strategies and their controls. Adaptive repair (AR) became the more advantageous the higher the initial cell density, compared with damage segregation (DS). At the start, 4, 8, 16 or 32 cells were randomly placed on the surface. Results are shown using log biomass ratios to make the horizontal line at log(ratio) = 0 (i.e., ratio = 1) a symmetry axis. The top row shows time courses for AR *versus* DS competitions, while the bottom row shows mean log(ratio)’s with standard deviations for all simulations, including controls, taken when biofilms had reached 154 µm (approximately 250 – 300 hours). Statistics for these competitions are shown in Table S1 and biofilm structures are plotted in Fig. 5 and File S1. Control time courses are shown in Fig. S7. Maximum specific growth rate was 1.2 h^-1^ and damage accumulation rate was set at 0.22 h^- 1^ and dependent upon specific growth rate for all strategies.

**TABLE 1.** All symbols, variables and parameters used

| Symbol | Name | Units |
| --- | --- | --- |
| $a$ | aging (damage accumulation) rate (constant) | $0.1 \text{ h}^{-1}$ |
| $a'$ | aging rate proportional to net specific growth rate | dimensionless,<br>0.22 |
| $\beta$ | investment in repair as a proportion of the total protein (fixed) | dimensionless,<br>0.07 |
| $\hat{\beta}$ | adaptive investment in repair as a proportion of the total protein | dimensionless,<br>between 0 and 1 |
| $\mu_{max}$ | maximum specific growth rate (Koch & Wang, 1982) | $1.2 \text{ h}^{-1}$ |
| $\mu(S)$ | specific growth rate as a function of substrate concentration | $\text{h}^{-1}$ |
| $K_S$ | Monod (half-saturation) constant (Koch & Wang, 1982) | $0.00234 \text{ g L}^{-1}$ |
| $P_{g,a}$ | protein invested into growth, active | fg |
| $P_{r,a}$ | protein invested into repair, active | fg |
| $P_{g,d}$ | protein invested into growth, damaged | fg |
| $P_{r,d}$ | protein invested into repair, damaged | fg |
| $P_{act}$ | active protein ($P_{g,a} + P_{r,a}$) | fg |
| $P_{dam}$ | damaged protein ($P_{g,d} + P_{r,d}$) | fg |
| $P_{gro}$ | protein invested into growth ($P_{g,a} + P_{g,d}$) | fg |
| $P_{rep}$ | protein invested into repair ($P_{r,a} + P_{r,d}$) | fg |
| $P_{tot}$ | total protein ($P_{g,a} + P_{r,a} + P_{g,d} + P_{r,d} = P_{act} + P_{dam}$) | fg |
| $P_{div}$ | threshold protein mass triggering division (46) | 620 fg (dry mass) |
| $Z$ | cellular age, or damaged protein as a proportion of total protein<br>($\frac{P_{dam}}{P_{tot}}$) | dimensionless,<br>between 0 and 1 |
| $\alpha$ | degree of asymmetry, or damage segregation | dimensionless,<br>between 0 (fully symmetric) and 1 (fully asymmetric) |
| $\theta$ | baby mass fraction; proportion of protein inherited by the new pole cell (normally distributed with mean 0.5 and standard deviation 0.025) | dimensionless,<br>between 0 and 1 |
| $Y_{\mu}$ | growth yield, the efficiency of converting glucose to active protein (Neijssel *et al.*, 1996) | $0.444 \text{ g g}^{-1}$ |
| $Y_r$ | repair yield, the efficiency of converting damaged protein to active protein (46) | $0.8 \text{ g g}^{-1}$<br>(assumed) |
| $r(P_{act}, P_{dam})$ | rate of repair | $\text{h}^{-1}$ |
| $S$ | substrate concentration | $\text{g L}^{-1}$ |
| $S_{in}$ | substrate concentration in the inflow (46) (used in the constant and chemostat environments) | 0.00234<br>(constant) or |
|  |  | 0.00324<br>(chemostat) g L<sup>-1</sup> |
| $S_{bulk}$ | substrate concentration in the bulk liquid (used in the biofilm environment) | 0.014222, 0.000889 or 0.003556 g L<sup>-1</sup> |
| $D$ | dilution rate (46) | 0.3 (chemostat) or 0.6 (biofilm bulk liquid) h<sup>-1</sup> |
| $\rho$ | biomass density (dry mass); assumed to be lower in biofilms due to the presence of extracellular polymeric substances (EPS). | 290 (constant and chemostat; (100)) or 201 (biofilm) g L<sup>-1</sup> (assumed) |
| $b_L$ | boundary layer thickness (unless otherwise stated) | 48 μm (assumed) |
| $\delta^2$ | ratio of maximum substrate transport to maximum substrate consumption rate | dimensionless, 0.0017, 0.0069 or 0.028 (assumed) |
| $\sigma_f$ | absolute deviation of biofilm front points from the mean front position; a measure of biofilm roughness | μm |
| $h_{max}$ | maximum biofilm thickness (unless otherwise stated), above which cells would be removed (here, the simulation was stopped when this height was reached) | 154 μm (assumed) |
| $D_G$ | diffusivity of growth substrate (glucose) | $6.7 \times 10^{-10}$ m<sup>2</sup> s<sup>-1</sup> (101) |
| $S_f$ | shove factor (102) | 1.10 (assumed) |

We decided to use the log biomass ratios as the best measure of fitness after comparing the performance of a range of different metrics when both strategies were identical (controls in Table S1 and Fig. S7). The ideal fitness measure should not be time-dependent, such that running simulations longer would not change outcomes. As the trends of the log biomass ratios in Figs. 6 and S7 show, using biomass at the end of the simulations (approximately 250 h) was appropriate since outcomes would not have changed had the simulations continued. Moreover, the fitness metric with its significance testing should never result in a statistically significant result when the two competitors were identical, as was the case with final growth rate (Table S1). Final growth rate was therefore not a suitable measure. Both biomass and population size could be used; they are strongly coupled, but the biomass changes are smoother than the cell numbers, hence, biomass was chosen as the fitness measure.

The biofilms formed from these equal mixtures of DS and AR, initially placed randomly on the substratum surface, show the spatial distribution of the strategies and the age and activity of the cells (Fig. 5 and S7). First, it can be seen that there was limited mixing of cells where neighboring ‘fingers’ touch (lines of different color standing out), consistent with our previous work (67, 68). This is due to the assumption that cells embedded in the biofilm matrix are not motile. Second, if finger-like cell clusters were similarly separated from other clusters, both strategies reached a similar height, suggesting that shrinking was negligible (Fig. 5; 4 cell initial density). This was despite repair leading to loss of material and cellular volume as growth could compensate for any losses when the rate of damage accumulation was proportional to growth rate. Third, the effect of increasing the initial cell density can also be seen. When there were only a few cells randomly placed on the substratum surface, it was likely that one cell happened to be further away from competing cells than any of the other cells. This lineage could therefore grow without much competition and became the highest finger producing the highest biomass, regardless of which strategy it followed. Hence, results at low cell density were more dependent on the random initial cell positions determining the distance to competitors than on competitiveness. The higher the initial density, the less influenced the results were by the stochastic initial attachment of cells on the surface. In these cases, AR had the advantage. Finally, comparing the distribution of young *vs* old cells with the distribution of active *vs* inactive cells (Fig. 5) shows that AR were all young throughout the biofilm, while DS created a mixture of young (rejuvenated) and old (damage-keeping) cells throughout the cell clusters, regardless of height. Note that cells are only active when they are at the top of their biofilm finger and when this finger is at least about as high as the neighboring fingers because only then do they receive sufficient substrate diffusing in from the bulk liquid, which is separated from the biofilm by a concentration boundary layer (63–65). Hence, whether a strategy still had cells in the active layer was determined by whether it still had cells growing close to the maximum biofilm height, and not by how many cells it had that were very young.

## DISCUSSION

Here, we investigated two alternative strategies for unicells to deal with intracellular damage: segregation or repair. Due to a trade-off between investing cellular resources into repair machinery *versus* growth machinery that is fundamental to all living organisms, repair is costly and therefore not obviously beneficial. This study is the first to compete these strategies in biofilms, representing spatially structured environments. We began by developing a new model for adaptive repair of cellular damage, whereby cells are able to sense and respond to current damage levels by investing into repair machinery accordingly, enabling cells to deal with spatial and temporal changes of conditions in biofilms. We found that in almost all conditions tested, adaptive repair was fitter than the asymmetric segregation of damage at division (Figs. 3 and 6). We also compared adaptive repair with our previous fixed optimal repair strategy (46) – in well-mixed environments where both strategies are suitable. Surprisingly, although cells with the adaptive repair mechanism had slightly lower growth rates than those with fixed optimal repair at the level of the individual (Fig. 2B), they outperformed them in the chemostat environment where there was competition for resources (Fig. 3). The likely reason for this is that with adaptive repair, there are more cells growing at the highest rate (Fig. 2C); as cells grow exponentially, any slight growth rate advantage will increase over time, resulting in a higher chance of division before stochastic removal from the chemostat.

When we initially applied the adaptive repair model to growth in biofilms we were surprised that the bases of biofilm ‘fingers’ of adaptive repairers were disappearing (Fig. 4). Since we assume that converting damaged into new material incurs a loss of 20% to account for the energy requirement of repair, the slowly growing cells at the base of the biofilm are not able to compensate for this 20% loss of biomass due to repair with new growth. Therefore, repairers shrink. This could be considered a biofilm specific disadvantage of repair. However, we think such extensive shrinking is unrealistic. First, to our knowledge, extensive shrinking of the volume of starving or dormant cells other than by cell division has not been observed and this may be due to the murein sacculus maintaining cell shape. Second, these biofilm structures with completely disappearing bases are not realistic as the slightest shear stress would detach these structures (66, 69, 70). Moreover, one would expect that the higher the rate of metabolism such as protein folding or respiratory electron transport, the higher the chance of damage arising such as protein misfolding or damage by reactive oxygen species (19, 71, 72). Indeed, organisms that grow more rapidly have been shown to also accumulate damage more rapidly (47, 73) and can have a higher rate of mortality (38). Therefore, we decided to make the simplest assumption that the rate of damage accumulation should be proportional to the specific growth rate of individual cells. In this case, biofilm fingers of adaptive repairers no longer had shrinking bases (Fig. 5) and now performed better than the damage segregators (Fig. 6 and Table S1), as may be expected when cell death is predominantly intrinsic rather than extrinsic, as in the constant and chemostat environments. Unfortunately, there are no empirical studies comparing the benefits of aging versus repair in biofilms, highlighting the dearth of studies of aging in the most common habitat of microorganisms.

The evolved extent to which unicells segregate damage asymmetrically varies substantially between species. This poses the question of whether this is due to differences in the mechanism of cell growth and division, differences in the ‘life span’ of their habitats or the degree to which these ‘unicells’ are actually multicellular with a clearer division of labor between germline and soma. The budding yeast, *Saccharomyces cerevisiae,* was the first unicell shown to age (21) and, in fact, is the only unicell for which the evidence of aging under benign conditions, rather than as a stress response, remains strong (9, 20, 32–38, 74–77). Its habitat is very rich in sugars but short-lived (78), and we argued previously (46) that investing resources into repairing the cell rather than reproduction is less advantageous when the habitat is (reliably) transient. However, the fission yeast *Schizosaccharomyces pombe* lives in the same kind of habitat (79). Also, the bacterium *Caulobacter crescentus* lives attached to surfaces that are decaying or consumed by zooplankton and therefore similarly transient, yet the evidence for senescence in *C. crescentus* has dwindled since our 2014 publication (9, 34, 80). This suggests that our previously proposed explanation – that morphologically asymmetric cell division and high external mortality due to short-lived habitats are necessary and sufficient conditions to see aging in unicells – needs to be revised as these conditions appear necessary but not sufficient. A third condition, nascent multicellularity, needs to be also met, see below.

Recent studies of the fission yeast, following single cells for many generations, provide clear evidence that asymmetric damage segregation does not occur under benign conditions. Instead, it appears to be a stress response to deal with misfolded proteins, which aggregate and then fuse into fewer and larger aggregates, facilitating the segregation of the damage into one aged daughter cell with a reduced growth rate and higher mortality (32, 33, 81). Spivey *et al.* (2017) (37) found that cell death in the fission yeast was actually not preceded by the characteristics of aging. Nakaoka & Wakamoto (2017) (38) also found no increase of mortality with age, instead, they found mortality to increase with growth rate. Moreover, they found that aggregates can get lost from old pole cells during division so they can rejuvenate.

Remarkably, oxidative stress reduced growth rate only transiently and protein aggregates present in the cell after stress did not affect growth. Apart from fission yeast, a recent study of *C. crescentus* by Iyer-Biswas *et al.* (2014) (34) found a trend of decreasing fecundity over generations only at 37°C, the highest temperature used. This could be considered a heat shock given typical lake temperatures where *C. crescentus* lives. Moreover, Schramm *et al.* (2019) (9) found no evidence for asymmetric segregation of protein aggregates, despite the morphologically asymmetric cell division in *C. crescentus*. In *E. coli*, the extent to which a cell segregates damage asymmetrically at division tends to increase with the severity of environmental stress that cells are exposed to (36). In *Mycobacterium tuberculosis*, segregation is critical for recovery from stress that resulted in damaged protein that cannot be repaired (35). An entirely different issue is survival during starvation, where aging has recently been proposed as an adaptive strategy based on the finding that starving *E. coli* cells, like non-starving humans, follow the Gompertz law of mortality (82). However, the Gompertz law has been shown to arise from a variety of processes, including tumor growth, growth of batch cultures and genetic or acquired susceptibilities to death (83–88), so no mechanistic conclusions can be drawn from finding that mortality follows the phenomenological Gompertz law.

Considering all evidence, the budding yeast appears to be the only studied ‘unicell’ where aging occurs in the absence of stress. However, neither its asymmetric cell division mechanism of budding, nor living in transient habitats, is sufficient to explain this difference. It seems to us that the common view of the budding yeast as a unicell may be mistaken and the missing ultimate reason why budding yeast ages is that it is, to some extent, a multicellular organism. First, the monophyletic yeast lineage (*Saccharomycotina*) is a branch of the *Ascomycota*, a division of fungi that have many mycelial forms with multicellular hyphae, so yeasts have a multicellular heritage (89). Second, yeast can easily evolve towards multicellularity when cluster formation is selected (30, 31). Third, while many lab strains are mutants in a gene required for filamentous growth, which would hamper genetic analysis, wild strains of yeast are dimorphic with a unicellular yeast and a pseudohyphal multicellular form under starvation (29). Thus, the ‘unicellular’ yeast is at the cusp of multicellularity. We propose that the combination of the budding mechanism of asymmetric growth and division, dispersal between transient habitats and nascent multicellularity are the ultimate reasons that the budding yeast is the exception that evolved aging as part of the normal life cycle rather than as a stress response.

Our study has several limitations. First, we have focused on the effect of damage on growth rate rather than mortality. This is partly for simplicity and partly because some studies show an increase of mortality with age (90) while others suggest mortality is random rather than increasing with age (37, 38, 79). Second, we neglect damage that had not been repaired before it became segregated or that would be prohibitively expensive to repair, since the work of Lin Chao’s group has covered this well. They showed that damage segregation under constant environmental conditions leads to separate steady state levels of damage in old and young lineage cells, meaning that growth rate and mortality of cells do not change over divisions (43, 44, 91–94). (Since there is no trend of deterioration and the replication of young and old lineages do not fit a soma germline distinction, it is probably better to refer to this as damage homeostasis rather than aging). Third, we avoided specific assumptions on mechanisms of damage repair or segregation that are organism specific as these are subject to evolution and our interest is the evolution of general strategies. Fourth, we have simplified the biofilm system to growth on flat, inert surfaces without detachment. While our rough, finger like biofilms capture typical aspects of biofilm structure, many processes and potential structures could not be covered in this study as the number of possible combinations is huge. Nonetheless, our study is the first to cover the extremes, from a perfectly mixed chemostat to a simple biofilm without any mixing (no motility of cells and only diffusive transport of substrate through boundary layer and biofilm). That the results were, surprisingly, essentially the same for both extremes, suggests that the findings hold for environments in between these extremes.

In conclusion, our model predictions are confirmed by the experimental literature showing that aging provides fitness advantages only if the following necessary and sufficient conditions are met in combination for a given organism: (i) presence of a cell division mechanism that clearly enables asymmetric division of all damage such as budding; (ii) predominant habitat is transient or the extrinsic mortality high for other reasons, favoring early reproduction; and a degree of multicellularity is present. Otherwise, aging is advantageous only as a stress response to deal with damage that failed to be repaired. Here, we have expanded the scope of this prediction substantially from previous work in constant and dynamic but spatially uniform environments by exploring biofilms as an exemplar of spatially structured systems thought to harbor the majority of microbes in the environment. In contrast to our original hypothesis, we found that repair is also better than damage segregation in biofilms.

## MATERIALS AND METHODS

We are following the standard ODD (Overview, Design concepts, and Details) protocol for describing individual-based models to facilitate comparison and review (Grimm et al., 2006, 2010).

### Purpose

The purpose of this study is to determine whether segregation or repair of damage is the optimal unicellular strategy for dealing with cellular damage in spatially structured systems such as biofilms. In order to do this, we expand upon our previous work in spatially uniform systems (constant and chemostat environments), Clegg *et al.* (2014) (46) by introducing a strategy for adaptive damage repair and further subdividing biomass into four rather than two components. All simulations described here were performed using the free and open-source modelling platform iDynoMiCS (individual-based Dynamics of Microbial Communities Simulator) (97).

### State variables and scales

#### Growth parameters

Growth parameters (Table 1) used are for *Escherichia coli*, where available, and are the same as those used by Clegg *et al.* (2014) (46). Note that ‘protein’ is representative of the whole biomass.

*Environments:* Three environments were used for this study: constant, chemostat and biofilm.

*Constant environment:* Substrate concentration is kept constant and population size is kept to 1,000 as new cells randomly replace existing cells.

*Chemostat environment:* The simulation domain behaves like a chemostat of size 1 µm^3^. A chemostat is a well-mixed system where fresh resources constantly flow in and cells and left- over resources constantly flow out, at the same dilution rate *D* (0.3 h^-1^; Table 1).

#### Biofilm environment

Only two dimensional simulations were used, to simplify analysis, however, the addition of a third dimension would not be expected to change results because both horizontal dimensions are equivalent (68). The domain size is (256 x 256) µm^2^, and the spatial grid for solving the diffusion reaction equation has a resolution of (4 x 4) µm^2^. The glucose concentration in the bulk liquid connected to the biofilm domain is kept constant throughout.

#### Length of time simulated

Constant and chemostat environment simulations were run for a maximum length of 500 days. Single species simulations were run for 500 days, and competition simulations were stopped earlier if only one species remained in the simulation domain. Some single-species, single-cell simulations were run for only three days when the purpose was to follow one old pole cell over sufficient generations. Biofilm simulations were run until a maximum biofilm height of 154 µm was reached.

### Process overview and scheduling

#### Growth, aging and repair

Cell growth is exponential, as growth rate is proportional to the current mass of the cell (but see about age dependence below). Cells divide once their total protein threshold, *P_div_* (Table 1), is reached (randomized by drawing from a normal distribution with given mean ± standard deviation; 0.5 ± 0.025). Cells are made of two types of biomass, referred to as protein: protein invested into growth machinery, or protein invested into repair machinery, and protein may be either active or damaged (Table 1). As cells grow, they make active protein that is invested into growth. Active protein is damaged in one of two ways: at a constant rate (*a*), as in our previous work (46), or at a rate that is proportional to the cellular specific growth rate (*a*′). If cells possess the ability to carry out repair, damaged protein is converted back to active protein, at a rate proportional to both the concentration of damaged protein and the concentration of active repair protein with rate constant *β*, but with an efficiency, or repair yield (*Y_r_*), of 80% (these processes are summarized in Fig. 1).

Individuals can differ in their strategy for dealing with cellular damage, but strategies are inherited and do not evolve. There are two cell division strategies: cells either divide symmetrically or asymmetrically. Here we only look at complete symmetry or asymmetry as partial asymmetry always gave intermediate results in our previous study (46). In an asymmetrically dividing cell, the ‘old pole’ daughter cell inherits all damage (up to capacity), while the ‘new pole’ daughter cell inherits none and is therefore rejuvenated. There are three repair strategies: (i) No repair; (ii) Investing into repair machinery at a fixed fraction (*β*) of newly formed biomass. The optimal fixed fraction of investment as a function of damage accumulation rate was previously determined for chemostats (46); and (iii) Adaptive investment into repair machinery (*β̂*) depending upon the current levels of damage within the cell. This adaptive repair strategy is new to studies that model aging and repair in unicells and was developed to account for the different specific growth rates of cells in a biofilm, which change in time and space. Altogether, there were six combinations of division and repair strategies used in this study (Fig. 1). All six combinations of division and repair strategies were used for initial, single-strategy investigations (*i.e.* Fig. 2), but only FR, AR and DS were used for competitions.

#### Mortality (intrinsic and extrinsic)

In all environments, cells may be considered dead when their age reaches 1, signifying that there is no longer any active protein within the cell (intrinsic mortality). Such ‘dead’ cells are assumed to remain physically intact and to continue to occupy space (only relevant for biofilms) since cell wall degradation is presumably a slow process (taking many residence times in the chemostat and longer than the times we simulate in biofilms).

*Constant environment:* a cell is removed at random each time a division occurs (extrinsic mortality).

*Chemostat environment:* cells are removed randomly with the outflow from the chemostat at the dilution rate (extrinsic mortality).

*Biofilm environment:* cells are not removed from the simulation for reasons given in the fitness section (no extrinsic mortality). Rather than removing the cells when the maximum biofilm height is reached (simulating detachment), we stop the simulation.

#### Biofilm structure

The formation of biofilm structure was investigated in the absence of aging and repair in order to find conditions that would give typical biofilm structures, *e.g.* a smooth, an intermediate and a rough biofilm. Earlier models have found that rougher and more finger-like biofilms tend to be produced when nutrient availability is limiting growth (Picioreanu et al., 1998; Dockery and Klapper, 2002; Olivera-Nappa et al., 2010). This can be achieved, *e.g.*, by reducing the substrate concentration in the bulk liquid *S_bulk_* or increasing the thickness of the boundary layer *b_L_* which both have a direct effect on the flux with which substrate diffuses into the biofilm, and therefore on the thickness of the actively growing layer (65). Dimensionless groups have previously been introduced to explain the combined effects of *S_bulk_* and *b_L_* and other parameters on biofilm structure (see the Supplementary Materials and Methods section for further explanation of these). Here we use:

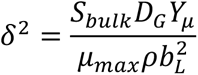

where *D_G_* is the diffusion coefficient of the growth substrate, *Y_μ_* is the growth yield, *μ_max_* is the maximal specific growth rate and *ρ* is the biomass density. In order to obtain biofilm structures that were smooth, intermediate or rough, *b_L_* was kept constant at 48 µm and *S_bulk_* took three different values (in g L^-1^): 0.014222 (smooth, *δ*^2^= 0.028), 0.003556 (intermediate, *δ*^2^= 0.0069), and 0.000889 (rough, *δ*^2^= 0.0017).

#### Time

Time is discrete. Since diffusion and biochemical reactions are on a faster timescale than growth and cell division, the diffusion-reaction equation is solved while the biomass distribution is fixed. Once the substrate concentration field is updated, it is kept fixed while the agents are stepped so they can grow and divide. Both diffusion-reaction and agent time steps were set to be the same at 0.01 h^-1^ in constant and chemostat environments, and 0.05 h^-1^ in biofilm environments where rates are lower.

#### Event scheduling

The simulation is entirely time stepped rather than event driven. The order in which agents are called in each time step is randomized.

#### Environments

Three environments were used for this study: constant, chemostat and biofilm.

*Constant environment:* Substrate concentration is kept constant and population size is kept to 1,000 as new cells randomly replace existing cells.

*Chemostat environment:* The simulation domain behaves like a chemostat of size 1 µm^3^. A chemostat is a well-mixed system where fresh resources constantly flow in and cells and left- over resources constantly flow out, at the same dilution rate *D* (0.3 h^-1^; Table 1).

Since the entire state of all agents and the environment is saved, simulations can be restarted with this output.

### Initialization

#### Environment

For the constant and chemostat environments, simulations are initiated with 1,000 cells (500 per strategy). Biofilm simulations are initiated with 4, 8, 16 or 32 cells (2, 4, 8 or 16, respectively, of each strategy), which are placed randomly on the substratum surface.

### Input

Almost all system and agent parameters are specified in an xml input file called ‘protocol’ file. Example input files can be found at https://github.com/R-Wright-1/iDynoMiCS_1.5.

### Sub-models

#### Mathematical skeleton

The following equations are for modeling growth, aging, and repair of individual cells. They are ordinary differential equations (ODEs). Their solution depends on conditions prescribed at one end of the interval of interest (Lick, 1989).

#### Individual Model Equations

The population is not modelled directly, but summary statistics are gathered and rates summed over all individuals. The substrate consumption rates of all individuals are gathered and summed and this total rate of substrate consumption enters the standard equation for chemostat substrate dynamics (+ inflow − outflow − consumption). Note that the net specific growth rate of an individual is also the sum of the rates of change for all four components of the cell. We give the differential equations for the change of the cell’s components below. Individuals do not have access to population level information and their behavior depends only on local conditions.

The biofilm environment consists of substrate concentration fields and a representation of the current biofilm structure (substratum surface, biofilm, biofilm boundary-layer interface and boundary-layer bulk-liquid interface). The environment is modelled as a continuum using partial differential equations (PDEs) to describe the diffusion of substrate and rates of substrate uptake (or product secretion) by the cells. For this purpose, the distributions of cellular masses and substrate consumption (reaction) rates are mapped to the grid used for solving the PDEs. The reaction diffusion PDEs are solved with a multigrid algorithm, see Lardon *et al.* (2011) (97) for more details.

Since adaptive repair has a variable fraction of repair machinery, the previously used *P_act_* and *P_dam_* have each to be split into two fractions:

> *P_g,a_*, *P_r,a_*, *P_g,d_* and *P_r,d_*, referring to growth machinery, active; repair machinery, active; growth machinery, damaged; and repair machinery, damaged, respectively (as in Table 1).

Thus, the total ‘protein’ of the cell (representing all biomass) is *P_tot_* = *P_g,d_* + *P_r,d_* + *P_g,a_* + *P_r,a_*.

We assume that damage is always toxic, *i.e.*, specific growth rate, due to some inhibitory effect of damaged material, decreases with the fraction of damaged protein, *Z*, equivalent to the age of the cell:

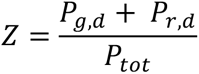

Toxic damage led to more pronounced differences between the strategies in our previous study, whilst not changing the fitness ranking of strategies apart from one case, at the lowest damage accumulation rate and only in the constant environment, where the differences between strategies were minute (46).

In cells of all strategies, growth of active protein depends on substrate concentration *S* following Monod kinetics:

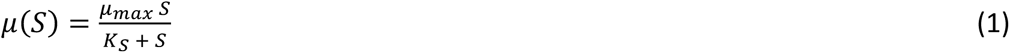

Where repair of damage takes place, the rate of repair is Michaelis-Menten like and proportional to damaged protein and active repair protein (see Clegg *et al.* (2014) (46) for further explanations):

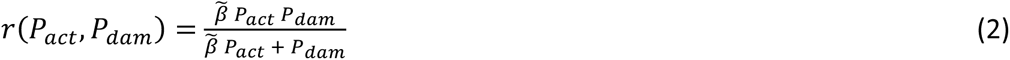

where *β̃ P_act_*represents the proportion of active biomass that is dedicated to repairing damaged biomass and a placeholder for the value actually used depending on the repair strategy. For fixed repair, it becomes the fixed fraction *β* of active protein *P_act_* that is repair machinery *β̃ P_act_*= *β P_act_*. For adaptive repair, it is replaced by the currently active repair machinery *β̃ P_act_* = *P_r,a_*, which is produced as a fraction of growth *β̂* calculated for each individual at every time step depending on its current fraction of damage (age *Z*) from the following equation:

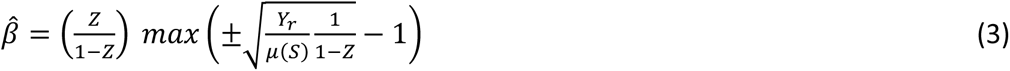

which is the value of *β* that maximizes the rate of active protein production and is derived from 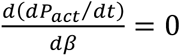.

For *μ*(*S*) in eq. 3, we do not take the gross specific growth rate according to eq. (1), but the net specific growth rate each individual cell calculates from its change of total mass from one iteration to the next, which due to inefficient repair could be less.

This gives the following differential equations for the four components of each individual cell for the case of toxic damage that is being repaired:

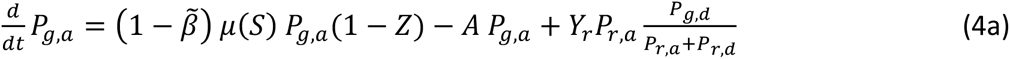

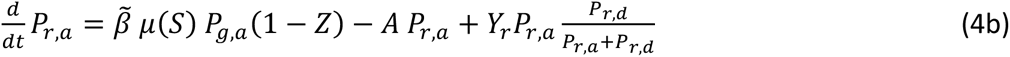

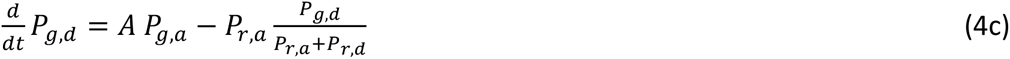

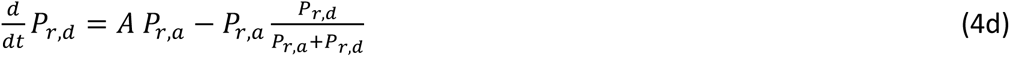

where *A* is a placeholder for the use of constant (*a*) or net specific growth rate proportional aging rate (*a*′):

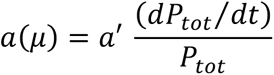

#### Cell division

Upon cell division, the post-division protein masses of the old pole cell are:

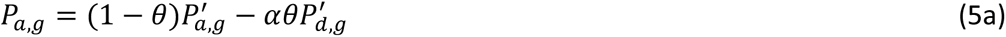

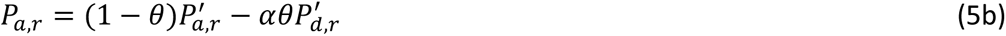

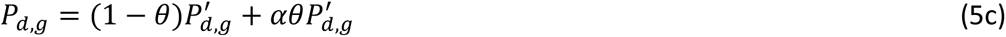

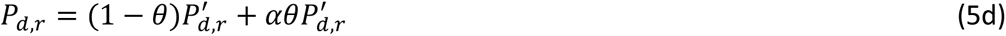

and those of the new pole cell are:

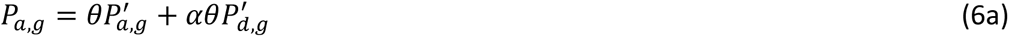

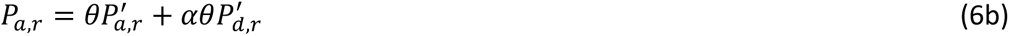

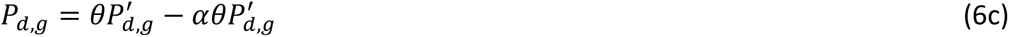

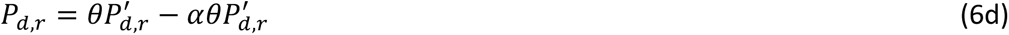

where the prime indicates the protein amounts in the pre-division cell, *θ* the proportion of protein inherited by the new pole cell and *α* the asymmetry of cell division (*α* = 1 fully asymmetric division, *α* = 0 fully symmetric division). However, if there is more damage than the old pole daughter cell can take (when 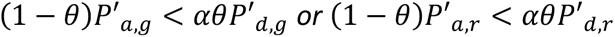 so the old pole-cell would inherit a negative quantity of active protein following eq. 5), the old pole cell is assumed to be filled with damaged protein:

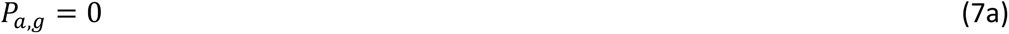

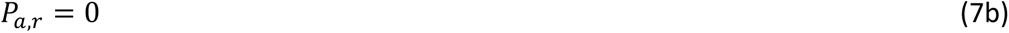

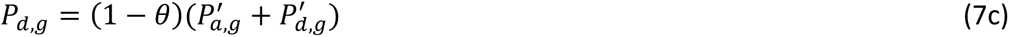

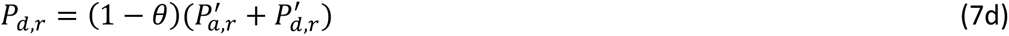

and the new pole cell inherits all of the active protein, plus the remainder of the damaged protein:

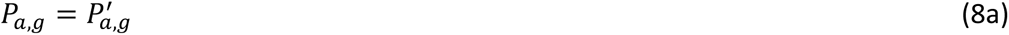

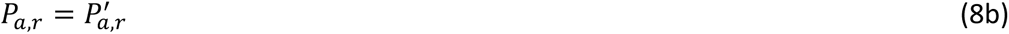

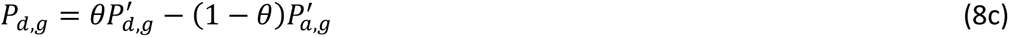

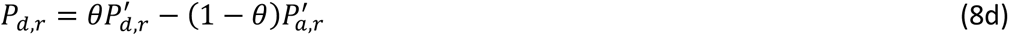

#### Aging

The growth independent aging rate of 0.1 h^-1^, used in Clegg et al. (2014), was suitable for constant and chemostat (spatially uniform steady state) environments, but was not suitable for biofilm (spatially structured) environments. Most cells within biofilms are not actively growing. Only those cells in the top or active layer of biofilms have access to substrate and are actively growing. It therefore makes sense to assume that non-growing cells that do not produce proteins or respire do not accumulate damage (19, 71, 72). Thus, the damage accumulation rate was assumed to be proportional to cellular specific growth rate.

In order to compare strategies across environments, we need to apply the same damage accumulation rate in all three environments. Hence, the new damage accumulation rate that is proportional to specific growth rate, *a*′, must be calculated to match the fixed damage accumulation rate in the spatially uniform environments (*a* = 0.1 h^-1^) where the specific growth rate is constant or predictable for a steady state. In the steady state of the chemostat, the net specific growth rate, µ_N_, is equal to the dilution rate, D, which was set to 0.3 h^-1^. In iDynoMiCS, the gross specific growth rate, *μ*_*G*_, is calculated with the Monod equation and depends on substrate concentration, S. However, how much the net specific growth rate is lower than the gross specific growth rate depends on the age of the cell, its rate of repair and its current level of investment into repair. It is therefore difficult to work out analytically so we had to run a number of simulations with different ratios of aging rates to specific growth rates to find that a value of 0.22 (dimensionless) would match the previously used constant aging rate for chemostats (Fig. S6).

#### Diagram of processes

A diagram containing a brief overview of all cellular processes is in Fig. 1.

### iDynoMiCS Simulations

#### Cell strategies

Six combinations of damage segregation and repair strategies were used in this study, but only the fittest three of these were used in biofilm simulations: asymmetric division without repair (DS), symmetric division with adaptive repair (AR), and symmetric division with fixed repair (FR). They are described in the schematic depicted in Fig. 1. Damage is considered to be toxic in all simulations as toxic damage leads to greater differences between strategies (46).

#### Comparison with previous fixed repair

In order to compare the new, adaptive repair strategy developed here with the previous fixed repair (as in (46), single strategy and competition simulations were run in iDynoMiCS. These simulations were initiated with 1,000 cells for single strategies or 500 of each strategy for competitions. The single strategy simulations were run for 3 days in the constant environment, where the ‘old pole’ cell was artificially kept in the simulation, rather than allowing random removal, to examine the consequences of adaptive repair on an individual cell (Fig. 2). Alternatively, they were run for 500 days, to examine the consequences of adaptive repair on populations of cells. Competitions between cells of different strategies were run in both the constant and chemostat environments for a maximum of 500 days, or until only one strategy remained in the simulation domain (*n*=10 for each).

#### Biofilm environment simulations

We initially carried out biofilm simulations with a single strategy without aging or repair to determine the parameters that would give rise to typical biofilm structures. For these, cells were initially placed evenly on the substratum surface. For all further simulations, with damage accumulation and repair, cells were initially placed at random on the substratum surface.

### Software and hardware used

iDynoMiCS 1.5 is free open source software written in Java (46, 97). Analysis scripts were written in Python 2.7.10 (Python Software Foundation, 2010). All source and analysis code can be found at https://github.com/R-Wright-1/iDynoMiCS_1.5.

## Supplemental Material

- Supplementary file containing Figures S1-S7, Table S1 and Supplementary Materials and Methods.
- **File S1.** Uploaded to Figshare: https://doi.org/10.6084/m9.figshare.11520534.v1. Includes further supplementary figures with biofilm plots for all biofilm simulations performed.

## Acknowledgements

RJW and TLRC were supported by BBSRC, UK; RW *via* a Midlands Integrative Biosciences Training Partnership PhD scholarship and TLRC *via* a University of Warwick Systems Biology Doctoral Training Centre PhD scholarship. RJC and JUK are grateful to the UK National Centre for the Replacement, Refinement & Reduction of Animals in Research (NC3Rs) for funding their development of individual-based models (IBMs) for the gut environment (eGUT grant NC/K000683/1). The funders had no role in study design, data collection and interpretation, or the decision to submit the work for publication.

